# *Drosophila* renal stem cells enhance fitness by delayed remodeling of adult Malpighian tubules

**DOI:** 10.1101/2021.12.03.471154

**Authors:** Chenhui Wang, Allan C. Spradling

## Abstract

*Drosophila* renal stem cells (RSCs) contradict the common expectation that stem cells maintain tissue homeostasis. RSCs are abundant, quiescent and confined to the peri-ureter region of the kidney-like Malpighian tubules (MTs). Although derived during pupation like intestinal stem cells, RSCs initially remodel the larval MTs only near the intestinal junction. However, following adult injury to the ureter by xanthine stones, RSCs remodel the damaged region in a similar manner. Thus, RSCs represent stem cells encoding a developmental redesign. The remodeled tubules have a larger luminal diameter and shorter brush border, changes linked to enhanced stone resistance. However, RSC-mediated modifications also raise salt sensitivity and reduce fecundity. Our results suggest that RSCs arose by arresting developmental progenitors to preserve larval physiology until a time in adulthood when it becomes advantageous to complete development by RSC activation.

**One-Sentence Summary:** Activated Drosophila renal stem cells rebuild the adult Malphigian tubules using a less efficient but more stone-resistant design.

---

Many tissues such as skin and intestinal epithelium have adult stem cells, which sustain homeostatic tissue function by regulated self-renewal and daughter cell differentiation. It is commonly assumed that all adult stem cells act to maintain the composition, morphology and function of their tissue in the face of fluctuating environmental conditions, stress and tissue damage (*1-3*). Here we show that the unusual properties of the renal stem cells (RSCs) found within the *Drosophila* adult Malpighian tubule (MT) suggest they represent a class of stem cells with an alternative function-tissue remodeling.

*Drosophila* MTs are functionally analogous to mammalian kidneys but with much simpler tissue structure (*4*). The MTs have little cell turnover under normal physiological condition but are vulnerable to damage under stress conditions (*5*). RSCs are only found within the peri-ureter region known as the stem cell zone (SCZ) of adult MTs (Fig. 1A), where they comprise 66% of tissue cells, but usually remain quiescent (*5*). The SCZ corresponds to the ureter and lower tubules which reabsorb 30% of fluid secreted by the upper tubules (*6*). The ureter is surrounded by visceral muscles which control fluid flow in the ureter via peristalsis (Fig. 1A) (*7*).

**Fig. 1.**
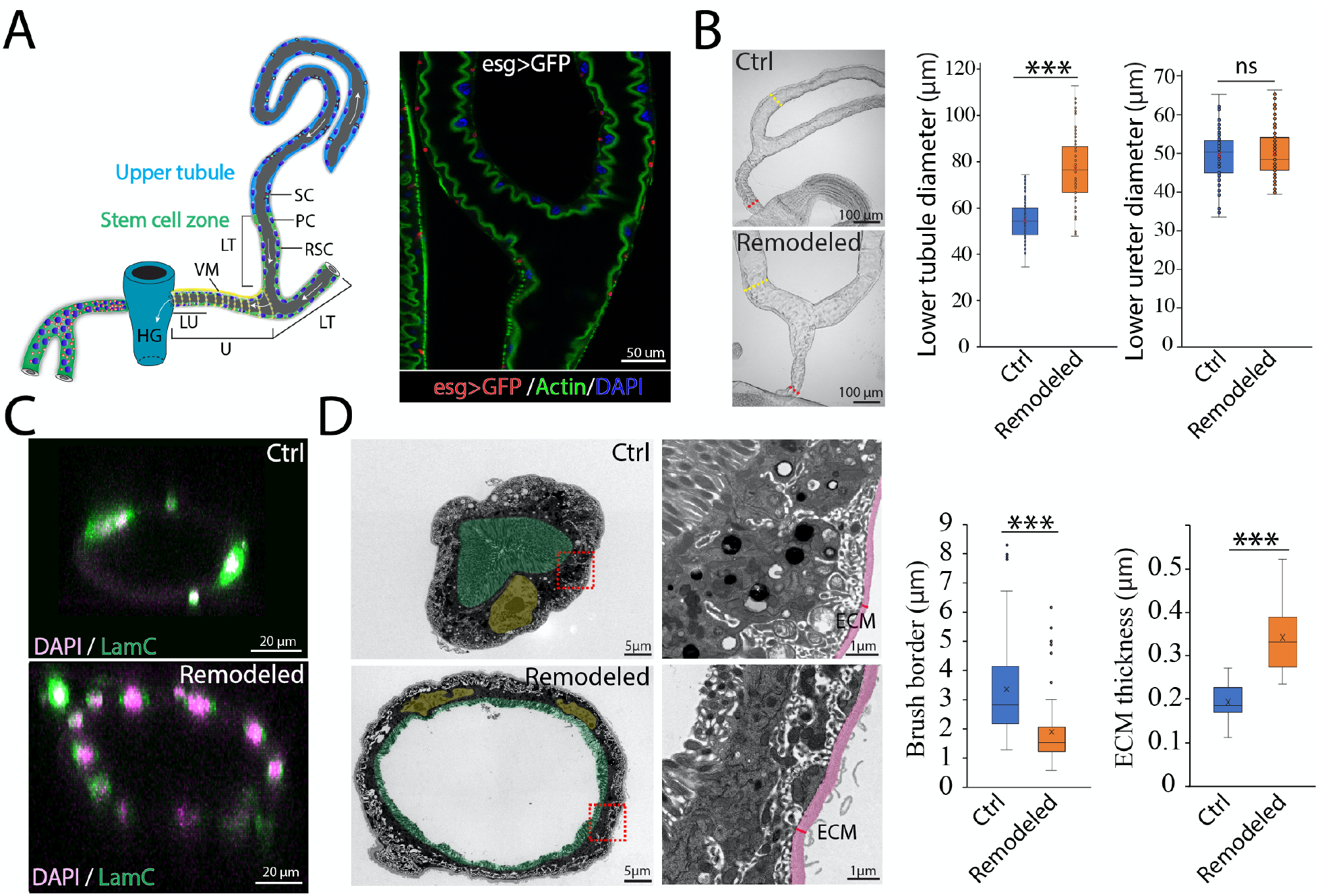
Morphological remodeling of adult Malpighian tubules. **(A)** Left: sagittal cross-section diagram of adult *Drosophila* MTs. Right: immunofluorescence (IF) of ureter and lower tubules of *esg-Gal4 >UAS-GFP* MT stained with phalloidin. RSC: renal stem cell, PC: principal cell, VM: visceral muscle, SC: stellate cell, LT: lower tubule, U: ureter, LU: lower ureter, HG: hindgut. White arrows: direction of fluid flow along the tubules toward hindgut. **(B)** RSC-mediated repair of the SCZ increases lower tubule diameter after 21 days at 18°C following genetic ablation of PCs at 29°C. Ctrl: *C507-Gal4*^*ts*^*/+* ; Remodeled: *C507-Gal4*^*ts*^ *>UAS-rpr+hid*. Quantitation at right. **(C)** IF of control (Ctrl) lower tubule and remodeled (Remodeled) lower tubule. **(D)** Pseudocolored TEM image showing thickened extracellular matrix (ECM, magenta) and shortened brush borders (green). PC nuclei (yellow). Quantitation at right. *** denotes two-tailed Student’s t test p<0.001, ns: not significant (p>0.05). Scale bars as indicated.

RSCs share a common origin with *Drosophila* intestinal stem cells (*8*). Both derive from the pool of adult midgut progenitors (AMPs), which extensively remodel the larval midgut yet leave the larval MTs almost intact except the lower ureter region where large larval principal cells (PCs) are replaced with smaller PCs (*9*). Adult RSCs barely replenish any tubule cells under normal conditions. However, RSCs can be activated when renal stones damage PCs located up to about ten cell diameters away. Intriguingly, instead of homeostatically replacing the damaged cells, RSC-derived replacement PCs are much smaller than preexisting PCs but more abundant, like PCs in the lower ureter region (*5*).

We genetically ablated adult PCs to more fully understand the nature of RSC-mediated lower tubule repair (see Supplementary Materials). While RSC-generated PCs are much smaller than preexisting principal cells, the overall DNA content of the tissue is largely restored (Fig. S1). However, we found that the diameter of lower tubules also increased significantly from 54.4 ± 8.2 µm (Mean ± SD, N=107 tubules) before repair to 77.0 ± 14.3 µm following RSC-mediated remodeling (p=1.79 × 10^−29^; N=80 tubules; Fig. 1B). In contrast, the diameter of the lower ureter, which undergoes RSC-mediated remodeling during pupal development, remained unchanged between control animals (49.5 ± 6.4 µm, N=75 ureters) and animals with remodeled MTs (50.0 ± 6.7 µm, N=64 ureters) (Fig. 1B). Despite their increased diameter, remodeled MTs remained a mono-layered epithelium albeit one containing more cells (Fig. 1C). Interestingly, the brush border of replacement PCs is significantly shorter than that of preexisting PCs (1.60 ± 0.75 µm versus 3.37 ± 1.88 µm, p=2.41× 10^−6^, N=8), whereas the extracellular matrix (ECM) thickness of the lower tubules substantially increases following repair (0.36 ± 0.09 µm versus 0.22 ± 0.05 µm, p=1.02× 10^−10^ N= 10) (Fig. 1D).

Since septate junctions (SJs) and adherens junction (AJ) are crucial for epithelial barrier function and cell adhesion in invertebrates, we examined the expression and localization of the conserved septate junction protein Coracle and the adherens junction core component Armadillo in lower tubules following regeneration (*10,11*). Coracle is localized to the apicolateral SJ of the preexisting principal cells, above the AJ domain to which Armadillo is localized, indicating the SJs form above the AJs in these ectodermally derived PCs (Fig. S2A-B). Following remodeling, localization of Coracle and Armadillo in replacement PCs remains the same (Fig. S2A-B). These results suggest RSC-mediated remodeling restores the barrier integrity of lower tubules in spite of the striking morphological alteration (Fig. S2C).

Kidney stones are a common renal disease in humans (*12*) that also affect insects such as *Drosophila* (*13*). The process of stone formation is readily studied using the *rosy (ry)* gene, which encodes xanthine dehydrogenase (XDH), and *ry* mutants develop xanthine-rich stones in their MTs at high frequency (*5,14*). We showed previously that xanthine stones elicit damage that triggers RSC-mediated repair in the SCZ (*5*). To further understand how xanthine stones form in *Drosophila* MTs, we first examined the distribution of stones in 3-7 day-old *ry* mutant animals that had been well fed after eclosion. Among *ry* mutant MTs bearing xanthine stones (N=211), 69.7% had visible stones exclusively in the SCZ, 22.3% had discernible stones in both the SCZ and the upper tubules, whereas only 8% exclusively had stones in the upper tubules (Fig. 2A-B). In addition, the vast majority of xanthine stone mass (94.7%, N=72) was found in the SCZ as assessed by area measurement (Fig. 2C). Therefore, RSCs are found in the MT region that is substantially more prone to acquire xanthine stones than other regions. This conclusion applied to both the anterior and posterior MT pairs, which differ in size and function. No differences in the average load of stones was found (Fig. 2D). Stone formation in anterior vs posterior tubules appeared to initiate independently, as little correlation in stone size was seen (Fig. S3A-D).

**Fig. 2.**
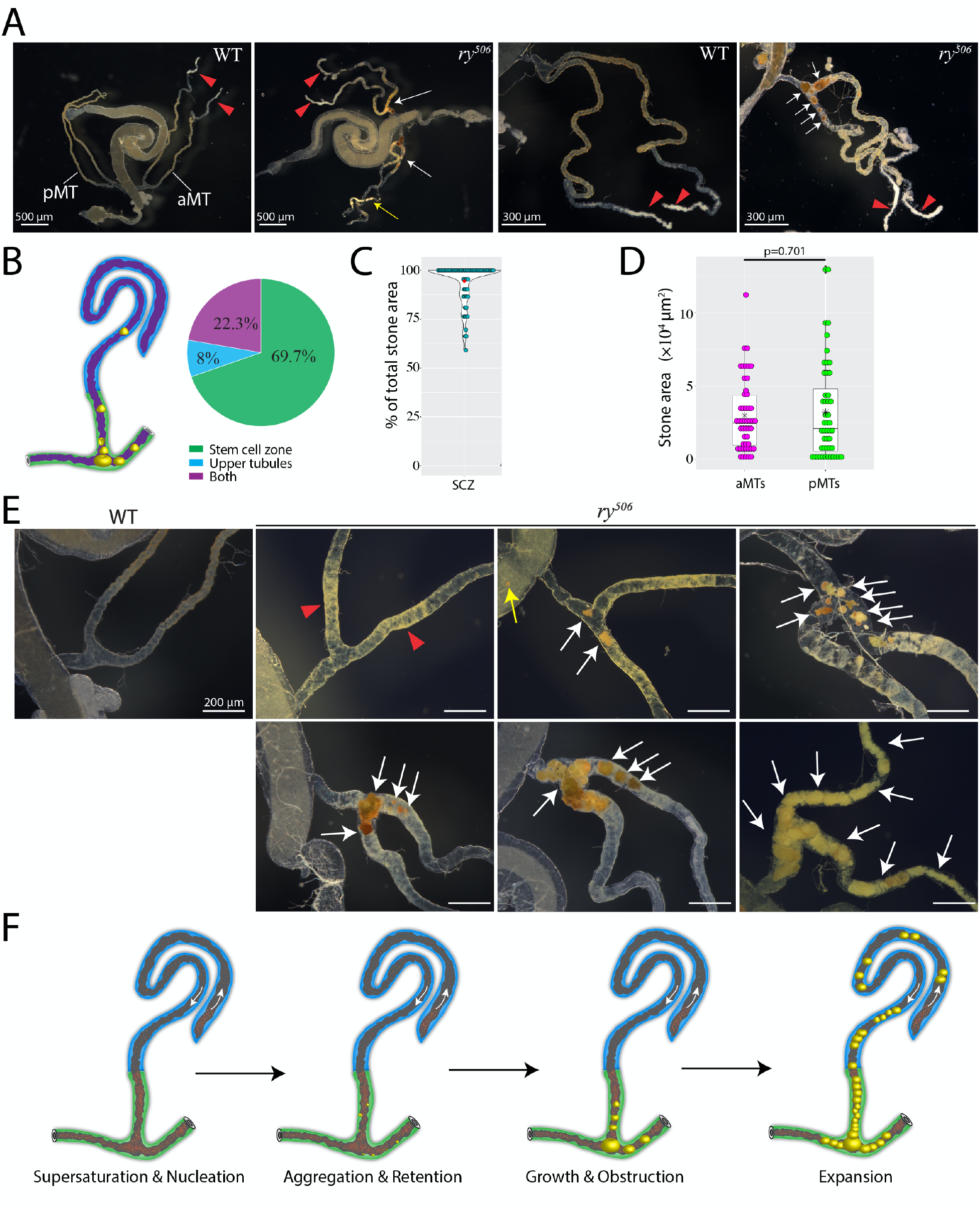
Progression of xanthine stone formation. **(A)** Preferential accumulation of xanthine stones (white arrows) in the ureter and lower tubules of *ry* mutants, or wildtype (WT) controls. Xanthine stones in upper tubule (yellow arrow). Red triangles denote calcium calculi in aMTs. Anterior MT pair (aMT); Posterior MT pair (pMT). **(B)** Stone distribution in MTs (3-7 day-old well-fed *ry* mutants. **(C)** Percentage of stone mass (area) in the SCZ versus upper tubules. **(D)** Stone mass (area) distributions in aMTs and pMTs. **(E)** Progression of xanthine stone formation (white arrows) in *ry* MTs. Red triangles denote lower tubules carrying tiny yellow particles. Stones are indicated in the hindgut (yellow arrow) and lower tubules (white arrows). Left panel shows control MT (WT). **(F)** Model of progressive xanthine stone formation in *ry* MTs. Scale bars 200 μm or as indicated.

Studying the early stages of stone formation revealed an additional connection between RSCs and stones. The lower tubules of *ry* mutants were more yellow in color compared to other MT regions or to wild type MTs, and small yellow particles could be seen to accumulate in the lower tubules prior to the appearance of stones (Fig. 2E). The lower tubule reabsorbs about 30% of fluid secreted by the upper tubules (*6*). Thus, the xanthine concentration in the lower tubules is more likely to become supersaturated than in the upper tubules, promoting stone nucleation (*15*). After nucleation, stones appeared sporadically in the SCZ, but were often found in the hindgut as well, indicating that small stones can be excreted into the hindgut (Fig. 2E). The dynamics of retention in the MTs and washout via the hindgut likely mediate stone progression, which can vary between tubules in the same animal (Fig. S3A-B). Once retained in MTs, stones gradually grow in size and aggregate with each other to form larger stones that lead to obstruction and RSC activation (Fig. 2E-F).

We previously found that RSCs enhance survival following Allopurinol treatment, which directly inhibits XDH (*5*). To investigate whether RSCs are evolutionarily conserved, we examined the MTs from adult *Drosophila pseudoobscura*, which is estimated to have diverged from *Drosophila melanogaster* about 33 million years ago (*16*). We found a population of diploid cells strongly resembling RSCs in their ureters and lower tubules (Fig. S4A). Moreover, Allopurinol-induced stones preferentially start to accumulate in this MT region in *Drosophila pseudoobscura*. RSCs can also repair the damage caused by stones in a similar manner as in *Drosophila melanogaster*: pre-existing principal cells are replaced by much smaller principal cells in stone-damaged tubules (Fig. S4B-C).

We carried out similar studies of a more distantly related fly, the lower Dipteran *Sciara coprophila*, which diverged from *Drosophila melanogaster* early in Dipteran evolution about 250 million years ago. In *Sciara*, the tubular epithelium only contains polyploid principal cells in the ureter and lower tubules, suggesting absence of RSCs (Fig. S4D). Adult MT remodeling may have evolved after the divergence of *Sciara* and *Drosophila*, or it may have been lost in *Sciara coprophila* since adults of this species are short lived compared to *Drosophila*.

MT remodeling by RSCs to increase tubule diameter and shorten brush border length may enhance stone resistance. To probe this possibility, we developed a scheme to rapidly induce xanthine stones by injecting flies twice with Allopurinol spaced by 24 hr (Fig. 3A). As we observed in *ry* mutants, stones induced by Allopurinol injection preferentially appeared in the lower tubules. After Allopurinol injection on four consecutive days, 89.9% ± 2.15% of control animals (N=90) carried discernible stones in at least one of the four MTs (Fig. 3B-C). In contrast, only 22.4% ± 10.5% of flies (N = 78) with remodeled MTs bore visible stones (Fig. 3B-C), a highly significant difference (p<0.001). In addition, the total stone area in affected animals was significantly decreased in remodeled compared to control MTs (5,840 ± 4,700 µm^2^ versus 12,900 ± 11,800 µm^2^, p=3.5×10^−5^) (Fig. 3d). Stones were more frequently observed in the hindguts of animals with remodeled MTs compared to control animals (Fig. 3B), consistent with increased expulsion of stones in remodeled MTs. A higher rate of loss likely contributes to the lower stone levels in remodeled MTs. Together, our data show that MT remodeling reduces the incidence of obstructing stones, and changes MT diameter and brush border length.

**Fig. 3.**
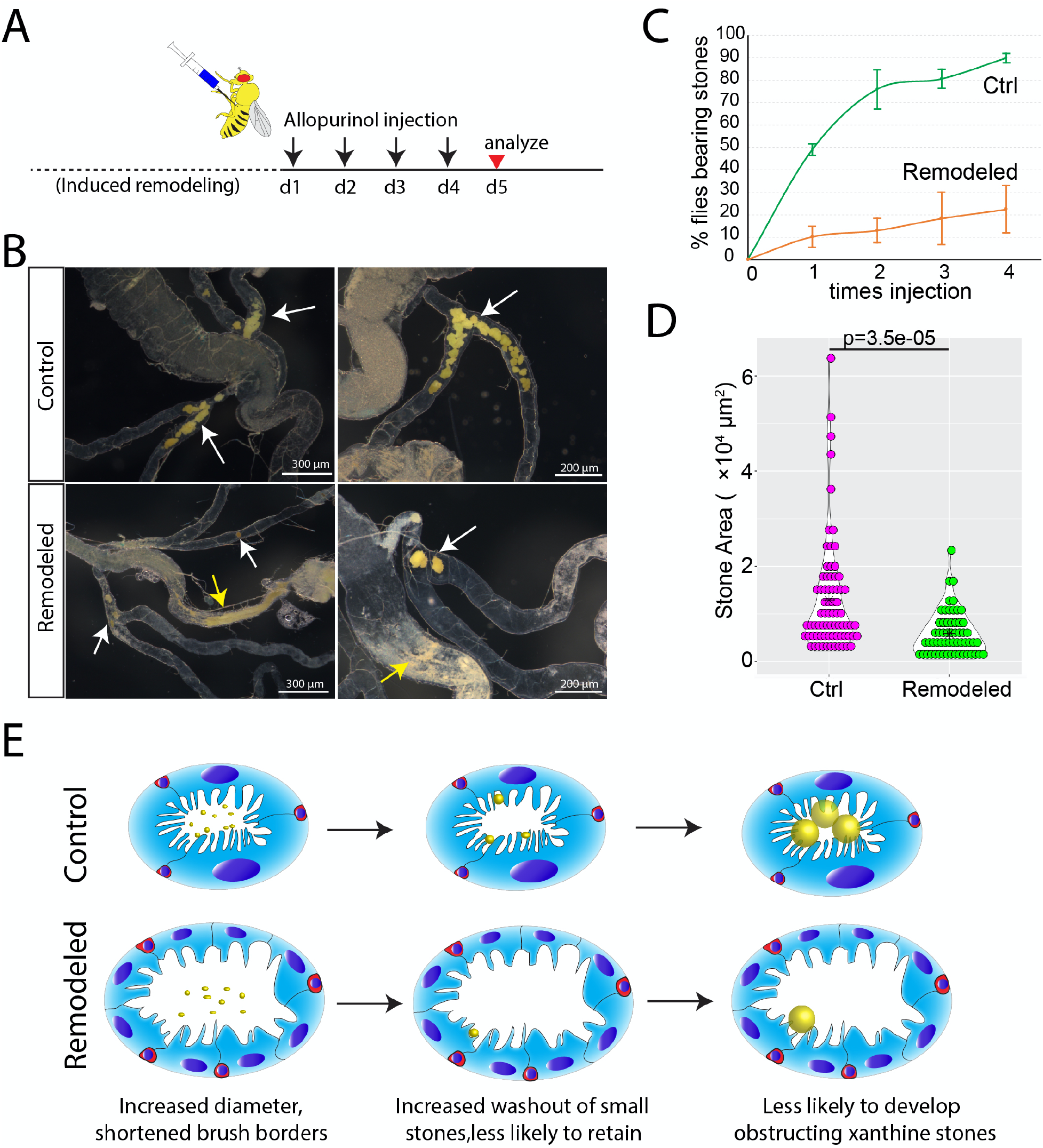
Remodeled MTs are stone-resistant. **(A)** Experimental scheme for xanthine stone induction by Alopurinol injection. d = day. **(B)** Xanthine stones (arrows) in control MTs (top) and remodeled (Remodeled) MTs (bottom). Stones in SCZ (white arrows); Stones in hindgut (yellow arrows). **(C)** Control (Ctrl) flies are more susceptible to induced stones than flies with remodeled MTs (Remodeled). Error bars denote SD. **(D)** Stone size (area) distributions in MTs from control (Ctrl) and in animals with remodeled MTs (Remodeled) after 4d of Allopurinol injection. **(E)** Model for the enhanced stone resistance following RSC-mediated remodeling of adult MTs, showing increased luminal diameter and shortened brush border of tubules within the SCZ, which increases stone washout into the hindgut. Polyploid PC nuclei (blue ovals); RSCs (red cells); stones (yellow spheres). Scale bars as indicated.

Since MT remodeling enhances stone resistance but does not take place as expected during pupal development, we looked for possible tradeoffs in fitness. We first examined whether MT remodeling affects lifespan. Interestingly, flies with remodeled MTs live as long and possibly longer than control animals fed on regular food at 18°C (Fig. 4A). We next examined whether adult MT remodeling affects fecundity. Eggs laid per female following induced remodeling of MTs were measured for six consecutive days and compared to control. From the fourth day on, female flies with remodeled MTs produced 23-27% fewer eggs per day (p< 0.05, N= 20 flies) (Fig. 4B). Decreased fecundity was not due to differences in the genetic background, as the number of laid eggs was comparable between *C507-Gal4*^*ts*^*/+* females and *C507-Gal4*^*ts*^*> UAS-rpr+hid* females without induced remodeling of Malpighian tubules (Fig. 4B). Together, these results indicate that RSC-mediated remodeling of adult MTs modestly reduces female fecundity.

**Fig. 4.**
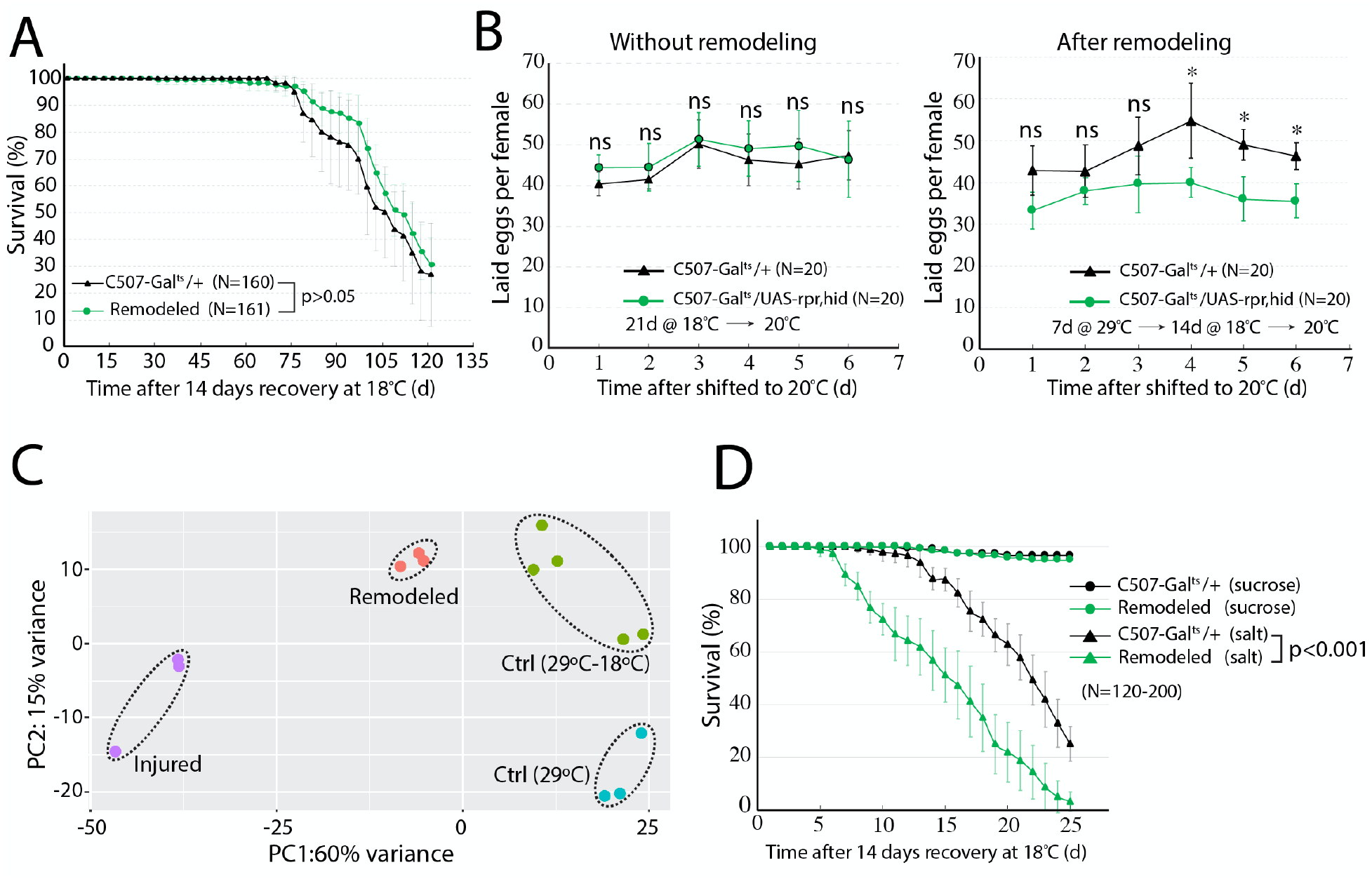
Remodeling of adult MTs compromises fecundity and high salt tolerance. **(A)** Survival curves (days) of controls (purple) and animals with remodeled MTs (green) at 18°C. **(B)** Fecundity (eggs laid per day per female) of control (*C507-Gal4*^*ts*^*/+*) or MT-remodeling competent (*C507-Gal4*^*ts*^*> UAS-rpr+hid*) females at 20°C. Left: Without remodeling (no heat treatment) before egg laying assay. Right: With MT remodeling (29°C for 7 days, 18°C for 14 days) before egg laying assay. **(C)** PCA plot of SCZ transcriptomes from Ctrl (*C507-Gal4*^*ts*^*/+*) females either at (29°C for 7 days; “29°C”), or at (29°C for 7 day, 18°C for 21 days; “29°C-18°C”); Injured and Remodeled (*C507-Gal4*^*ts*^*> UAS-rpr+hid*) females either at (29°C for 7 days; “Injured”), or (29°C for 7 day, 18°C for 21 days; “Remodeled”). Dots within dashed ellipses = biological replicates. **(D)** Survival curves of flies with control (c507-Gal^ts^/+) or remodeled (Remodeled) MTs fed on 5% sucrose with or without 250 mM NaCl; “salt”. Error bars indicate SD, * denotes p < 0.05, ns: not significant with two-tailed Student’s t-test.

To further assess the differences between the remodeled renal tubules and uninjured renal tubules, we compared the transcriptomes of the SCZ before and after remodeling using mRNA sequencing. Principal component analysis (PCA) and hierarchical clustering based on global gene expression showed that control and remodeled lower tubules express similar transcriptomes (Fig. 4C, Fig. S5A), suggesting they support very similar physiological functions. Nevertheless, the transcriptomes were not identical. 652 genes were differentially expressed with fold change > 2 and padj < 0.05, between remodeled and control MTs using DESeq2 (*17*). Among these genes, 360 genes were upregulated whereas 292 genes were downregulated in remodeled MTs compared to control. We subsequently performed GO enrichment analysis using Metascape (*18*). Interestingly, ECM-receptor interaction (dmel04512) stood out as the top enriched GO term among the upregulated genes. ECM genes including *vkg, Col4a1, LanB1, LanA, troll, wb, tig, tsp, ppn* were all upregulated after remodeling, agreeing with our previous observation of a thickened ECM following remodeling (Fig. 5B-C). These changes may be needed to maintain mechanical stability in light of the increased tubule diameter. In addition, a group of genes involved in transmembrane transport including sodium ion transport and potassium ion transport were upregulated, making the “GO:0055085 transmembrane transport” the second most enriched GO term among upregulated genes (Fig. 5B-C).

Given that the expression of transmembrane transport genes is already elevated in remodeled MTs under normal condition, we reasoned that these tubules might be more sensitive to salt-stress. Consequently, we modified a previously reported salt-stress assay to examine the salt tolerance capacity of flies bearing wild type MTs or remodeled MTs (*19*). The survival of flies bearing wild type MTs or remodeled MTs was comparable when fed a 5% sucrose-only diet. In contrast, flies bearing remodeled MTs died substantially faster than control animals when subjected to a high salt-stress diet containing 250 mM NaCl mixed with 5% sucrose (Fig. 4D). These results suggest that remodeling of adult MTs compromises their salt tolerance, possibly because an increased tubule diameter changes the surface to volume ratio in an unfavorable manner for processes like salt resistance that are highly dependent on surface-localized transporters.

Previous studies of stem cells that maintain tissues with active cell turnover suggest that they function to maintain tissue homeostasis in the face of cell loss, environmentally mediated damage, and fluctuating resources (reviewed in *1-3, 20-22*). By adjusting rates of cell production, proportions of different downstream cell types, and engaging in competition with genetically distinct cells for retention in a niche, the essential structure and function of many tissues can be maintained by homeostatic stem cells throughout life. However, our experiments show that RSCs represent a different class of stem cells. RSC-mediated repair of adult *Drosophila* lower tubules doesn’t restore tissue morphology and homeostasis, but mediates an irreversible tissue makeover, that takes place only once during adulthood. If the effects were uniformly positive, then RSCs would likely resemble other developmental progenitors and carry out beneficial remodeling prior to adulthood, or during major tissue regeneration. However, we found that the RSC-mediated makeover can exert either positive and negative fitness effects depending on circumstances beyond organismal control.

RSCs appear to be developmental progenitors that evolution has transformed into stem cells that persist into adulthood. This is similar to the proposed origin of female germline stem cells from primordial germ cells that become stabilized in a niche rather than further developing into germ cells (*23*). However, RSCs are progenitors stabilized by becoming quiescent, because this maximizes the ability to control when their morphological remodeling occurs. In many instances this would occur after reproduction has largely taken place, but before kidney stones have had time to form. We suggest this class of stem cells be called “makeover stem cells,” and suggest they are fairly common. However, makeover stem cells are much harder to identify than homeostatic stem cells, because they will remain quiescent under most circumstances, resembling differentiated tissue cells, like RSCs. Only under the appropriate conditions will they manifest their stem cell character.

Mammalian kidneys may possess makeover stem cells that are similar to RSCs. Our studies highlight the striking similarities in the process of renal stone formation between *Drosophila* and mammals (*13,14*). Indeed, loss of *XDH* generates frequent xanthine stones in both *Drosophila* and human excretory systems. Renal stones often first strike the *Drosophila* lower tubule, which acts to concentrate the tubular fluid and controls fluid flow via constriction of encircling ureter muscles. The mammalian counterparts of the *Drosophila* lower tubule with analogous functions are the renal papillae and renal pelvis (*24*). Interestingly, Randall’s plaques which are believed to act as nidi for urinary stone formation are predominantly attached to renal papillae protruding into the renal pelvis (*25*), like the MT brush border. Quiescent kidney papillary label-retaining cells have been reported to be able to regenerate medullar tubules upon severe kidney injury in mice (*26,27*). It would be interesting to learn whether medullar tubule regeneration increases stone resistance in mammals. Deepening our understanding of the cellular and molecular mechanisms underlying renal stone formation and cellular responses via makeover stem cells such as RSCs and possible mammalian counterparts, will advance our ability to prevent and treat kidney stones, a growing medical and economic burden globally (*12*).

## ACKNOWLEDGEMENTS

We are grateful to John Urban for providing *Sciara coprophila*, Ethan Greenblatt for providing *Drosophila pseudoobscura*, the Bloomington *Drosophila* Stock Center and Developmental Studies Hybridoma Bank for *Drosophila* strains and reagents. We thank Allison Pinder and Frederick Tan for RNA sequencing. We are grateful to Mike Sepanski for assistance with electron microscopy.

## Author contributions

Conceptualization: C.W., A.C.S.; Methodology: C.W., A.C.S.; Investigation: C.W.; Visualization: C.W., A.C.S.; Funding acquisition: A.C.S.; Project administration: A.C.S; Supervision: A.C.S; Writing – original draft: C.W., A.C.S.; Writing – review & editing: C.W., A.C.S.

## Data availability

Bulk mRNA-seq data reported here have been deposited at NCBI under the accession number: PRJNA772753

## Supplementary Materials

Materials and Methods

Figs. S1 to S5

