## Supplemental Methods and Figures for "*Drosophila* renal stem cells enhance fitness by delayed remodeling of adult Malpighian tubules"

### Materials and Methods

##### *Drosophila* stocks and husbandry

*Drosophila* stocks are reared on standard cornmeal molasses yeast food and maintained at room temperature ( 23-25°C) unless otherwise specified. A dash of dry yeast is routinely sprinkled on the surface of cornmeal molasses yeast food before use unless otherwise specified. The following stocks were used in this study: $ry^{506}$ (BDSC#4405), *OregonR* (BDSC#25211), *C507-Gal4* (BDSC#30840), *tub-Gal80*$^{ts}$ (BDSC#7019) UAS-RFP (BDSC#32222), *esg-Gal4;tub-Gal80*$^{ts}$*,UAS-GFP(28), UAS-rpr,UAS-hid (29).*

##### Immunostaining and microscopy

Malpighian tubules left attached to the gut were dissected in Grace's insect medium or 1×PBS. Samples were then fixed in 4% paraformaldehyde in PBS on a nutator for 20 min. After washing three times with PBT (1×PBS + 0.1%TritonX-100) for 5 min each, samples were blocked with 5% NGS in PBT for 1 h followed by incubating with primary antibody at 4°C overnight. Samples were washed three times with PBT and then incubated in secondary antibody for 2-3 hours at room temperature or 4°C overnight. After 3× 5 min washes with PBT, samples were counterstained with 100 ng/ml DAPI (and 1:1000 fluorophore-conjugated phalloidin if needed) for 5 min at room temperature. Samples were then washed twice with PBT for 5 min each and mounted in 50% glycerol. The antibodies used in this study: mouse anti-Lamin C ( 1:10, Developmental Studies Hybridoma Bank, Cat # LC28.26), mouse anti-Cora (1:50, DSHB Cat # C566.9 ), mouse anti-Arm (1:5, DSHB Cat # N27A1 ) , mouse anti-Cut (1:10, DSHB Cat # 2B10), Alexa 568 goat anti-Mouse (1:300, Thermo Fisher Cat # A11004), Alexa488 goat anti-

Mouse (1:300, Thermo Fisher Cat # A11001).Fluorescence images were taken with a Leica TCS SP8 confocal microscope.

**Induced remodeling of the SCZ**

We induced remodeling of the SCZ using genetic ablation of preexisting principal cells as previously described (*5*). Briefly, *tub-Gal80$^{ts}$;C507-Gal4,UAS-RFP* (referred to as *C507-Gal4$^{ts}$* for brevity) female flies were crossed to *UAS-rpr,UAS-hid* animals to produce *C507-Gal4$^{ts}$> UAS-rpr,hid* flies at 18$^{o}$C. 3-5 day old *C507-Gal4$^{ts}$> UAS-rpr,hid* flies were first shifted to 29 $^{o}$C for 7 days ( "Injured" ) and then shifted back to 18$^{o}$C for at least 14 days to allow completion of remodeling. *C507-Gal4$^{ts}$* female flies were crossed to *OregonR* males to produce *C507-Gal4$^{ts}$/+* which were used as control.

**Total volume of nuclei in the SCZ**

The wild type SCZ was marked by *C507-Gal4 >RFP*. The regenerated SCZ can be easily discerned as the replacement principal cells are much smaller compared to those in the upper tubules. z-stack images were acquired using a Leica TCS SP8 confocal microscope with a 20X objective lens (NA=0.75). The SCZ was selected to make 3D surface reconstruction of nuclei based on DAPI staining. The volume of surface masked SCZ was calculated by IMARIS v9.2.1 (Bitplane, RRID: SCR_007370) .

**Flies for RNAseq**

C507-Gal4$^{ts}$ animals were crossed to *OregonR* or *UAS-rpr, UAS-hid* animals at 18$^{o}$C. 3-5 day old *C507-Gal4$^{ts}$/+* females and *C507-Gal4$^{ts}$> UAS-rpr+hid* females were selected from above crosses and were subjected to temperature shift paradigm. After shifted to 29$^{o}$C for 7 days, half of the *C507-Gal4$^{ts}$/+* females and *C507-Gal4$^{ts}$> UAS-rpr+hid* females were dissected to collect the ureters and lower tubules. The samples were referred to as " Ctrl (29$^{o}$C) " and " Injured ", respectively. The rest of *C507-Gal4$^{ts}$/+* females and *C507-Gal4$^{ts}$> UAS-rpr+hid* females were shifted back to 18°C for 21 days prior to dissecting the ureters and lower tubules. These samples were referred to as " Ctrl (29°C - 18°C) "and " Remodeled ", respectively. About 100 flies were dissected in nuclease-free PBS on ice for each replicate. The samples were stored in 500 µl of TRIzol (Invitrogen 15596026) and stored at -80°C until all samples were ready for RNA extraction.

**mRNA sequencing**

The RNAs were then extracted following manufacturer's instructions. cDNA libraries were processed as previously described (*5*). 75bp single-end reads were aligned to the *Drosophila melanogaster* genome (dm6) using HISAT2 2.1.0 (*30*). Read counts per gene were calculated using htseq-count (*31*). Differentially expressed genes were identified using DESeq2 (*17*).

**Electron microscopy**

Samples from flies with desired genotypes were dissected and processed as previously described (*5*).

**Measurement of brush border and ECM**

The brush border length and ECM thickness were measured using EM images at 4,000× or 20,000× magnification from at least 8 different tubules of indicated genotypes, respectively. For each EM image, 5-8 subregions were randomly selected to be measured in Fiji (*32*) .

**Egg laying**

For egg laying assays, females of appropriate genotypes were subjected to the temperature shift in order to induce remodeling of MTs as described above. *OregonR* males were provided to mate with females at 1:1 ratio throughout the temperature shift. After 14 days at 18°C, males were replaced with young *OregonR* males. Twenty pairs of males and females of each genotype were subsequently divided into four groups and transferred to four separate bottles containing molasses plate with wet yeast at 20°C. The molasses plates were changed every 24 hours and the eggs laid on the plates were scored every 24 hours for six consecutive days.

**Survival under high salt-stress condition**

For salt feeding, a previous reported assay was modified by adding 1% melted agarose to the liquid diet to make solid food (*19*). Control flies and flies with remodeled MTs were switched from normal food to the agarose based diets containing 5% sucrose only (as control) or 5% sucrose + 250 mM NaCl mixed at 18°C. Flies were examined for survival daily and transferred to fresh food every other day.

**Allopurinol injection**

Flies with appropriate genotypes were injected with about 100 nl of injection solution (3mM Allopurinol mixed with 10 mg/ml Blue No.1 in 1× PBS and then transferred to regular corn meal food containing 3mM Allopurinol. A dash of dry yeast was sprinkled on food surface before use. Flies were injected for four consecutive days with an interval of 24 hours. Survival of the injections on all days was more than 90%.

**Quantification of stone area**

*Drosophila* MTs left attached to the gut were dissected in Grace's insect medium or 1×PBS. MTs were then imaged with a Nikon SMZ1500 stereo microscope equipped with HR Plan Apo 1× WD 54 objective, HR Plan Apo 1× WD 54 objective and Infinity 3 Lumenera camera within 30 min after dissection. Stones were manually outlined using the freehand selection tool in Fiji and the area was acquired using Fiji (*32*).

**Statistical analysis**

Student's t-test was used to determine the statistical significance in most cases except for the survival rate comparison, for which log-rank test was used. ns denotes $p > 0.05$, * denotes $p < 0.05$, ** denotes $p < 0.01$, *** denotes $p < 0.001$.

**Data availability**

Data has been uploaded to BioProject at NCBI under the accession code PRJNA772753.

(reviewer link:

https://dataview.ncbi.nlm.nih.gov/object/PRJNA772753?reviewer=7nbe6kpo7cbcp5o3etjjvgj9e0

)

Supplementary Figures

**Fig. S1 The total DNA content of the SCZ can be restored after remodeling. (A)** Representative micrographs of the SCZ from control and regenerated MTs. The 3D surface rendering of the nuclei in the SCZ was based on DAPI staining using IMARIS software. **(B)** The overall volume of nuclei is comparable between control and regenerated SCZ.

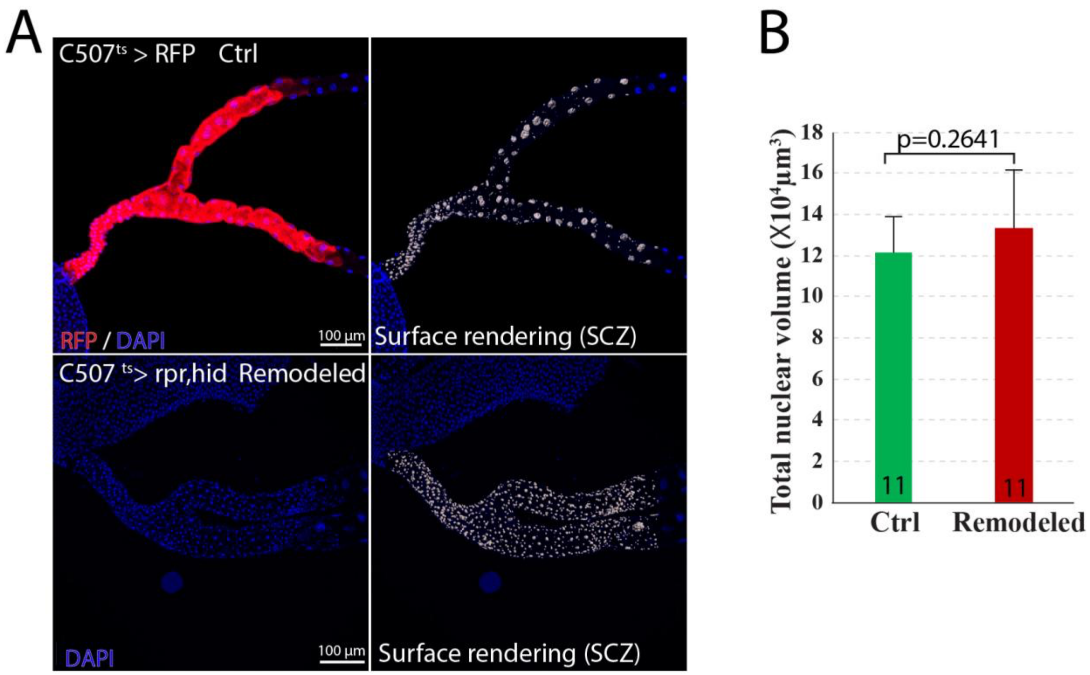


**Fig. S2 | The barrier integrity of Malpighian tubules can be restored following repair. (A)** Counterstaining of F-actin cytoskeleton and septate junction component Cora in control and regenerated SCZ. Note the apicolateral localization of Cora (denoted by white triangles) in preexisting principal cells as well as regenerated principal cells. **(B)** Staining of the adherens junction protein Arm in the wild type and regenerated SCZ. Note the basolateral localization of Arm (denoted by white triangles) in preexisting principal cells as well as regenerated principal cells. **(C)** A drawing depicting the morphological remodeling of adult lower tubules upon regeneration.

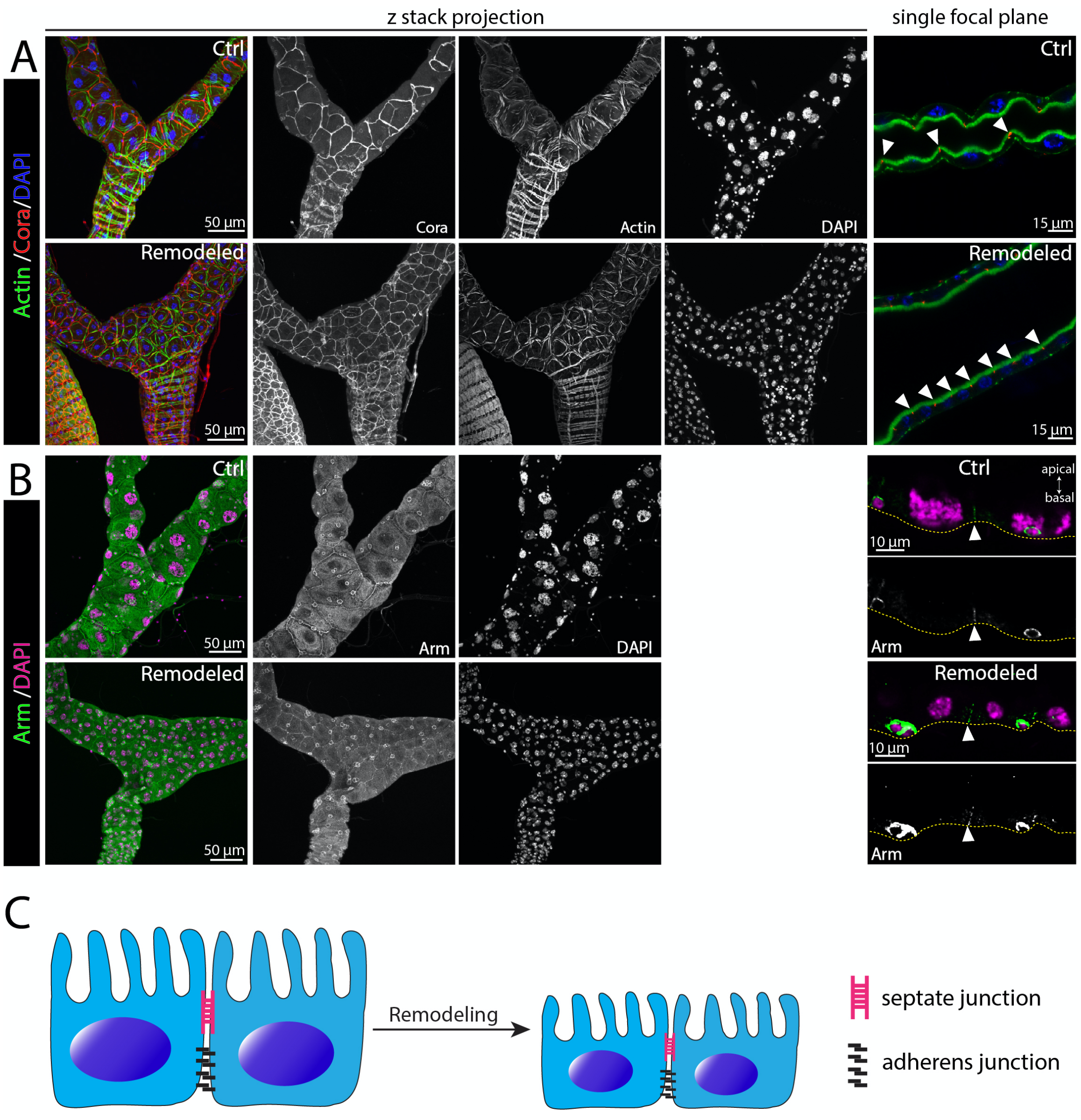

**Fig. S3  Obstructing xanthine stones caused by loss of *ry*.  (A-D)** Micrographs showing xanthine stones in anterior pair (denoted by white arrows) and posterior pair (denoted by yellow arrows) of MTs from *ry* mutants.  Xanthine stone formation in aMTs and in pMTs  are seemingly independent events.

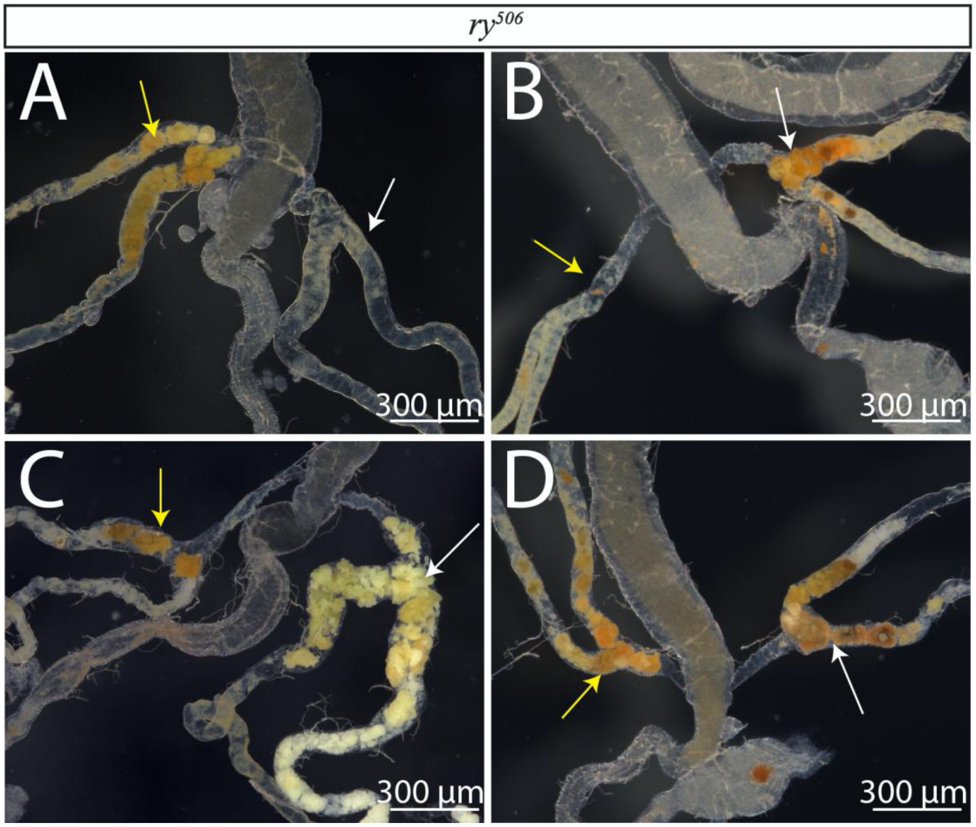


**Fig. S4 |  Conserved mode of RSC-mediated repair of adult MTs between *Drosophila melanogaster* and *Drosophila pseudoobscura*. (A)** Immunofluorescence micrograph of the ureter and lower tubules of an adult female wildtype *Drosophila pseudoobscura*  stained with 2B10 antibody (anti-Cut)  under normal condition.  Yellow arrows denote diploid cells that are presumably renal stem cells. **(B)** Representative image of the ureter and lower tubules of *Drosophila pseudoobscura* carrying Allopurinol-induced xanthine stones (red triangles). Note the supernumerary cells adjacent to xanthine stones. **(C)** Quantification of nuclear volume of preexisting PCs and replacement PCs in the SCZ from animals without or with Allopurinol treatment, respectively. Bar denotes the average value. *** denotes p<0.001 with Student's t test.

**(D)** Representative image of the ureter and lower tubules of an adult female wildtype *Sciara coprophila* stained with phalloidin. Note the absence of diploid cell population in the tubule epithelium.

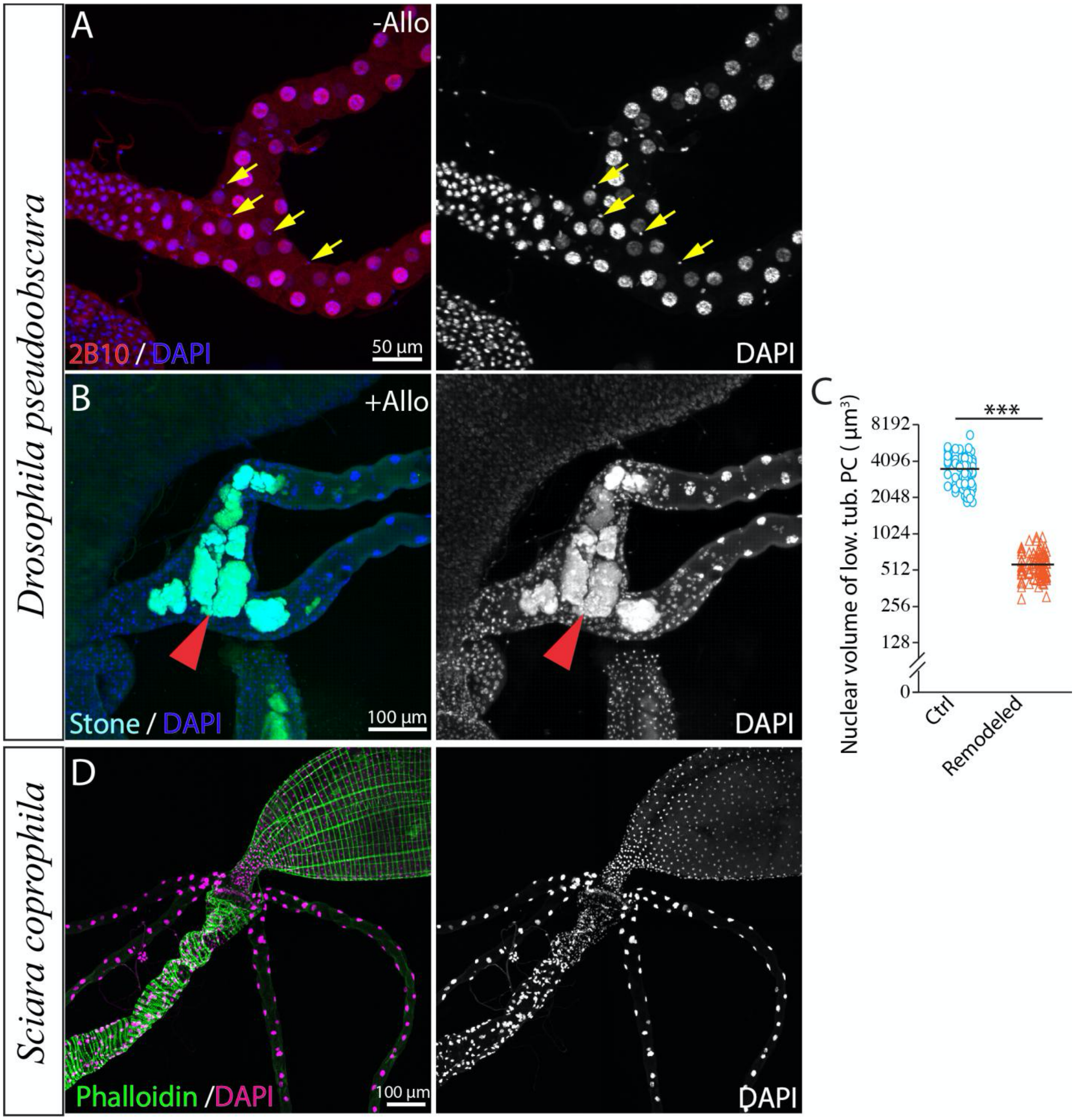

**Fig. S5 | Transcriptome comparison between the SCZ of remodeled MTs and that of control MTs. (A)** Heatmap showing the hierarchical clustering of samples based on global gene expression. **(B)** Left: Volcano plot showing the differentially expressed genes between the transcriptomes of Remodeled and Ctrl (29°C - 18°C) groups. Differentially expressed genes (padj<0.05, FoldChange >2) were identified using DESeq2. Upregulated genes and downregulated genes after remodeling are shown in red and blue dots, respectively. Right: GO enrichment of upregulated and downregulated genes respectively. GO enrichment analysis was performed using Metascape (*18*). **(C)** Heatmap showing the genes involved in ECM-receptor interaction (upper) and transmembrane transport (lower) that were upregulated in MTs after remodeling compared with control, respectively.

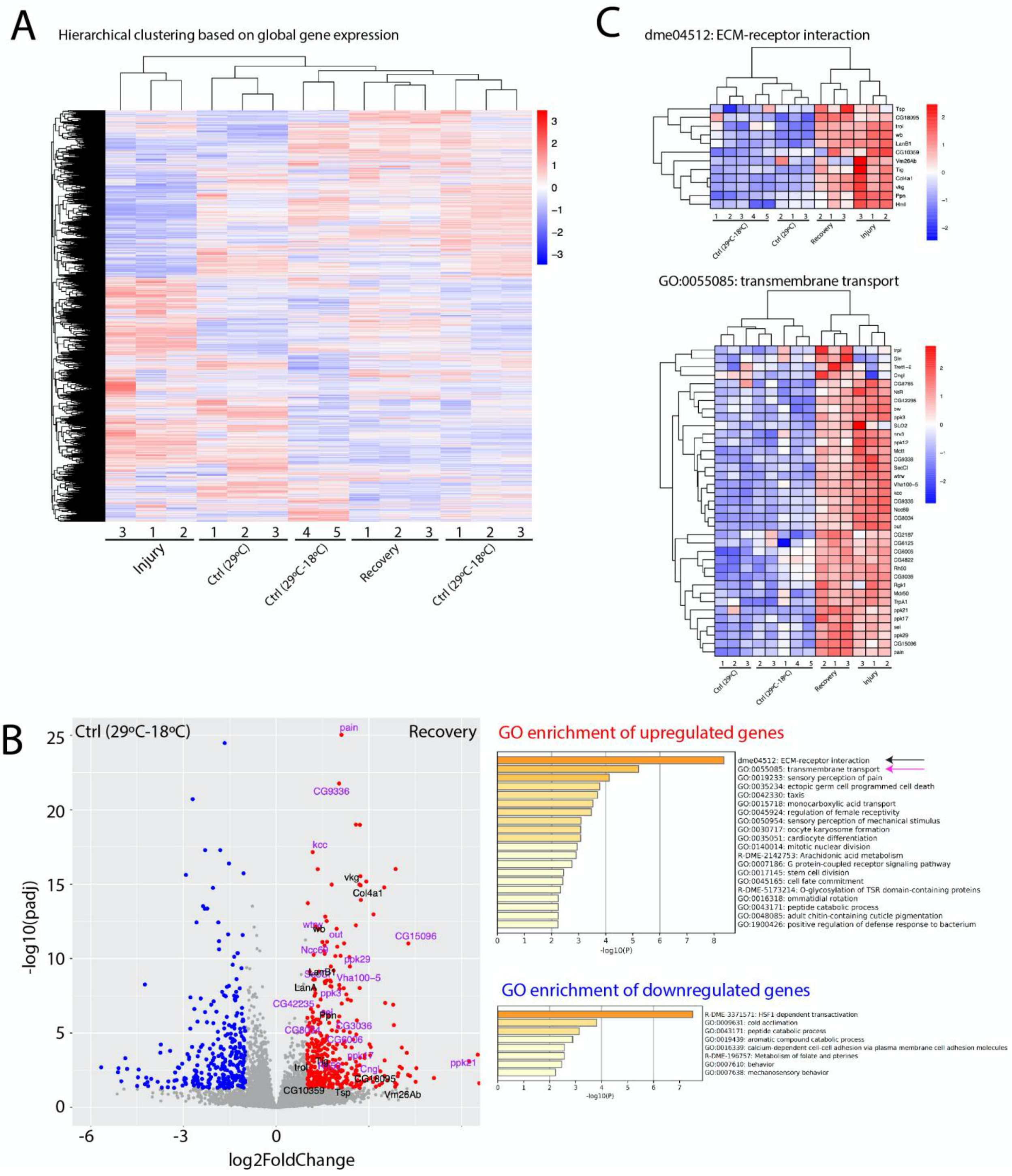

A
Hierarchical clustering based on global gene expression
Injury
Ctrl (29°C)
Ctrl (29°C-18°C)
Recovery
Ctrl (29°C-18°C)
B
Ctrl (29°C-18°C)
Recovery
-log10(padj)
log2FoldChange
GO enrichment of upregulated genes
dme04512: ECM-receptor interaction
GO:0055085: transmembrane transport
GO:0019233: sensory perception of pain
GO:0035234: ectopic germ cell programmed cell death
GO:0042330: taxis
GO:0015718: monocarboxylic acid transport
GO:0045924: regulation of female receptivity
GO:0050954: sensory perception of mechanical stimulus
GO:0030717: oocyte karyosome formation
GO:0035051: cardiocyte differentiation
GO:0140014: mitotic nuclear division
R-DME-2142753: Arachidonic acid metabolism
GO:0007186: G protein-coupled receptor signaling pathway
GO:0017145: stem cell division
GO:0045165: cell fate commitment
R-DME-5173214: O-glycosylation of TSR domain-containing proteins
GO:0016318: ommatidial rotation
GO:0043171: peptide catabolic process
GO:0048085: adult chitin-containing cuticle pigmentation
GO:1900426: positive regulation of defense response to bacterium
-log10(P)
GO enrichment of downregulated genes
R-DME-3371571: HSF1-dependent transactivation
GO:0009631: cold acclimation
GO:0043171: peptide catabolic process
GO:0019439: aromatic compound catabolic process
GO:0016339: calcium-dependent cell-cell adhesion via plasma membrane cell adhesion molecules
R-DME-196757: Metabolism of folate and pterines
GO:0007610: behavior
GO:0007638: mechanosensory behavior
-log10(P)
C
dme04512: ECM-receptor interaction
Ctrl (29°C-18°C)
Ctrl (29°C)
Recovery
Injury
GO:0055085: transmembrane transport
Ctrl (29°C)
Ctrl (29°C-18°C)
Recovery
Injury
